# Hepatic MCPIP1 protein levels are reduced in NAFLD patients and are predominantly expressed in cholangiocytes and liver endothelium

**DOI:** 10.1101/2022.08.16.504094

**Authors:** Natalia Pydyn, Justyna Kadluczka, Piotr Major, Tomasz Hutsch, Kinga Wołoszyn, Piotr Malczak, Dorota Radkowiak, Andrzej Budzynski, Katarzyna Miekus, Jolanta Jura, Jerzy Kotlinowski

**Author notes:** equal contribution as first author (Natalia Pydyn, Justyna Kadluczka). corresponding author, Jerzy Kotlinowski, Address: Gronostajowa Street 7, 30-387 Krakow, Poland. **Ethics approval statement**, All human tissue samples were collected according to the established protocol approved by the Local Bioethics Committee (approval no. 122.6120.263.2016). **Patient consent statement**, All subjects provided consent for the study.

## Abstract

Nonalcoholic fatty liver disease (NAFLD) is characterized by the excessive accumulation of fat in hepatocytes. NAFLD can range from simple steatosis to the aggressive form called nonalcoholic steatohepatitis (NASH), which is characterized by both fatty liver and liver inflammation. Without proper treatment, NAFLD may further progress to lifethreatening complications, such as fibrosis, cirrhosis or liver failure. Monocyte chemoattractant protein-induced protein 1 (MCPIP1, alias Regnase 1) is a negative regulator of inflammation, acting through the cleavage of transcripts coding for proinflammatory cytokines and the inhibition of NFκB activity.

In this study, we investigated MCPIP1 expression in the liver and peripheral blood mononuclear cells (PBMCs) collected from a cohort of 36 control and NAFLD patients hospitalized due to bariatric surgery or primary inguinal hernia laparoscopic repair. Based on liver histology data (H&E and Oil Red O staining), 12 patients were classified into the nonalcoholic fatty liver (NAFL) group, 19 into the NASH group and 5 into the control (non-NAFLD) group. Biochemical characterization of patient plasma was followed by expression analysis of genes regulating inflammation and lipid metabolism. The MCPIP1 protein level was reduced in the livers of NAFL and NASH patients in comparison to non-NAFLD control individuals. Additionally, in all groups of patients, immunohistochemical staining showed that the expression of MCPIP1 was higher in the portal fields and bile ducts in comparison to the liver parenchyma and central vein. The liver MCPIP1 protein level negatively correlated with hepatic steatosis but not with patient BMI or any other analyte. The MCPIP1 level in PBMCs did not differ between NAFLD patients and control patients. Similarly, in patients’ PBMCs there were no differences in the expression of genes regulating β-oxidation (*ACOX1*, *CPT1A*, and *ACC1*) and inflammation (*TNF*, *IL1B*, *IL6*, *IL8*, *IL10*, and *CCL2*), or transcription factors controlling metabolism (*FAS*, *LCN2*, *CEBPB*, *SREBP1*, *PPARA*, and *PPARG*).

We have demonstrated that MCPIP1 protein levels are reduced in NAFLD patients, but further research is needed to investigate the specific role of MCPIP1 in NAFL initiation and the transition to NASH.

## Introduction

Nonalcoholic fatty liver disease (NAFLD) is defined as the accumulation of excessive fat in the liver in the absence of excessive alcohol consumption and a lack of any secondary cause. Lipid accumulation in hepatocytes is a hallmark of the first stage of this disease and is called nonalcoholic fatty liver (NAFL) [1]. Although NAFL remission can be relatively easily achieved by lifestyle modifications and dietary changes, in approximately 25% of patients, the disease progresses to nonalcoholic steatohepatitis, hepatic fibrosis, liver cirrhosis and hepatoma [2]. In addition to fat accumulation, the activation of liver-resident macrophages, together with the recruitment of monocytes, neutrophils and lymphocytes into the liver parenchyma, is another hallmark of NASH. Both features have been associated with ongoing hepatic inflammation and NAFLD progression [3].

The immune system plays a critical role in NAFLD progression; thus, the analysis of proteins that regulate inflammation may shed new light on the NAFL-to-NASH transition. One protein that negatively regulates the inflammatory reaction is monocyte chemoattractant protein-induced protein 1 (MCPIP1). MCPIP1 is an endoribonuclease that binds to the 3’UTR fragments of mRNA and digests stem-loop structures. This endoribonuclease activity of MCPIP1 shortens the half-life of selected transcripts and therefore reduces the amount of proteins expression [4]. Moreover, MCPIP1 is responsible for the degradation of translationally active transcripts and is particularly important in the initial stage of inflammation [5]. It was also shown that MCPIP1 inhibits maturation of pre-miRNAs by digestion hairpin structures, which leads to a reduction in the miRNA cellular pool [6]. The anti-inflammatory properties of MCPIP1 have also been confirmed *in vivo*. Mcpip1 knockout mice spontaneously develop a systemic inflammatory response that leads to splenomegaly, lymphadenopathy, and hyperimmunoglobulinemia and ultimately leads to death within 12 weeks [7].

Mcpip1 acts as an important regulator of liver homeostasis both in mice fed chow and mice fed a high-fat diet [8]. Additionally, its deletion in murine liver epithelial cells recapitulates the features of primary biliary cholangitis, which manifests as excessive proliferation of intrahepatic bile ducts, bile duct destruction, inflammatory infiltration and fibrosis [9]. Sun and coworkers described a protective role of Mcpip1 in liver recovery after ischemia/reperfusion injury. Mcpip1 ameliorates liver damage, reduces inflammation, prevents cell death, and promotes tissue regeneration [10].

Despite data from mouse models, the functions of MCPIP1 in the human liver are still not known. Thus, in the current study, the MCPIP1 protein level was analyzed in a cohort of 36 patients who were divided into non-NAFLD, NAFL or NASH groups based on liver histological analysis according to NAFLD activity score (NAS) grading. For the first time, we demonstrated the diminished expression of MCPIP1 in the livers of NAFL and NASH patients in comparison to non-NAFLD control individuals. Additionally, the analysis of MCPIP1 distribution in liver tissue by immunohistochemical staining showed lower levels of MCPIP1 expression in the parenchyma and central vein than in the portal fields and bile ducts.

## Materials and methods

### Collection of blood and liver samples

Samples were collected from 36 patients (24 female and 12 male, BMI ranged from 36 to 70) who underwent bariatric surgery in The Second Department of General Surgery, Jagiellonian University Medical College (Krakow, Poland). Exclusion criteria included HCV/HBV/HIV infection, autoimmune diseases, cancer and alcohol abuse. Out of 36 patients, 15 were diabetic. Patients who were hospitalized due to primary inguinal hernia and qualified for laparoscopic transabdominal preperitoneal repair were enrolled in the non-NAFLD group. Blood for the analysis of blood count, plasma biochemistry and isolation of PBMCs was collected from fasting patients before bariatric surgery. Liver samples were acquired during bariatric surgery. One liver sample was placed in formalin (for histological analysis). The second liver sample was snap-frozen in liquid nitrogen and then stored at −80 °C (for protein analysis). All human tissue samples were collected according to the established protocol approved by the Local Bioethics Committee (approval no. 122.6120.263.2016).

### Blood analysis

All blood tests were measured routinely on the day of admission to the hospital in University Hospital laboratories with ISO 9001 certificates, using comparable laboratory methods.

### Isolation of peripheral blood mononuclear cells

Ten milliliters of EDTA-anticoagulated blood was collected from each patient and transferred to 15 ml tubes. In the first stage, blood was centrifuged at 400 x g for 10 minutes, and the plasma was removed. Then, the remaining blood was replenished with PBS to a volume of 10 ml and layered on top of 5 ml of Ficoll solution (1.077 g/ml). Gradients were centrifuged at 400□×□*g* for 30 min at room temperature in a swinging-bucket rotor without the brake applied. The layer of mononuclear cells was collected and washed twice with PBS by centrifugation. Finally, the cells were counted and suspended in appropriate buffers for RNA or protein extraction. Approximately 10^7^ cells were obtained from each patient.

### Protein isolation and western blot

PBMCs from patients were lysed using RIPA buffer (25 mM Tris-HCl, pH 7.6; 150 mM NaCl; 1% sodium deoxycholate; 0.1% SDS) supplemented with Complete Protease Inhibitor Cocktail (Roche) and PhosSTOP Phosphatase Inhibitor Cocktail (Roche). A bicinchoninic acid assay was used to assess the protein concentrations. Liver samples from patients were lysed in whole cell lysis bufer (62.5 mM Tris–HCl, pH 6.8; 2% SDS; 25% glycerol; 5%β-mercaptoethanol) supplemented with Complete Protease Inhibitor Cocktail (Roche) and PhosSTOP Phosphatase Inhibitor Cocktail (Roche). NanoDrop 1000 Spectrophotometer (Thermo Fisher Scientific) was used to assess the protein concentrations. Then, 50 μg of protein was separated on a 10% SDS-PAGE polyacrylamide gel. After wet transfer onto PVDF membranes (Millipore), the membranes were blocked in 5% skim milk and then incubated with primary antibodies at 4 °C overnight. On the following day, the membranes were washed and incubated with secondary antibody for 1 h at room temperature. Chemiluminescence was detected after 5 min of incubation with ECL™ Select Western Blotting Detection Reagent (GE Healthcare) in a ChemiDoc chemiluminescence detector (BioRad). The following antibodies were used: rabbit anti-MCPIP1 (1:1000; GeneTex), mouse anti-β-actm (1:4000; Sigma), rabbit anti-p-IGF1R^T_yr_1135/1136^/IRTyr^1150/1151^ (1:1000; Cell Signaling), rabbit anti-IR (1:1000; Cell Signaling), anti-p-Akt^Ser473^ (1:1000; Cell Signaling), rabbit anti-IGF1R (1:1000; Cell Signalling), rabbit anti-Akt (1:1000 Cell Signalling), rabbit anti-PPARγ (1:1000; Cell Signaling), rabbit anti-PPARα (1:1000 GeneTex), peroxidase-conjugated anti-rabbit (1:30000; Cell Signaling) and peroxidase-conjugated anti-mouse (1:20000; BD).

### RNA isolation and real-time PCR

Total RNA from PBMCs was isolated using Fenozol (A&A Biotechnology). A NanoDrop 1000 Spectrophotometer (Thermo Fisher Scientific) was used to assess the RNA concentration and quality. For reverse transcription, 1 μg of total RNA, oligo(dT) primer (Promega) and M-MLV reverse transcriptase (Promega) were used. Real-time PCR was carried out using Sybr Green Master Mix (A&A Biotechnology) and QuantStudio Real-Time PCR System (Applied Biosystems). Gene expression was normalized to elongation factor-2 (EF2), and then, the relative transcript level was quantified by the 2^deltaCt method. The primer sequences (Genomed/Sigma) and annealing temperatures are listed in Table S1 in the Supplementary Material.

### Histological analysis

Livers were fixed in 4% buffered formalin. Afterwards, tissue fragments were divided into two parts. One piece was immersed in a 30% sucrose solution overnight for cryoprotection and then frozen in Tissue-Tek^®^ OCT medium at −80 °C. Frozen sections were stained using the Oil Red-O (ORO) method for fat deposition and photographed under 100× magnification. Ten images of each section were randomly obtained and subsequently analyzed by the Columbus Image Data Storage and Analysis System (Perkin Elmer) with an algorithm adapted for Oil Red-O stained sections. The second piece was processed using standard paraffin procedures, and 5-μm paraffin tissue sections were stained with hematoxylin and eosin (H&E) and then visualized using a standard light microscope (Olympus BX51; Olympus Corporation, Tokyo, Japan). Immunohistochemical MCPIP1 staining was performed using primary anti-MCPIP1 antibody (1:200, Genetex) and EnVision Detection Systems Peroxidase/DAB, Rabbit/Mouse (Dako, Denmark).

Histopathological analyzes were performed by two independent experimental pathologists in a blinded way. Histopathological assessments and microphoto documentation were made using a Axiolab A5 light microscope with Axiocam 208 color and ZEN 3.0 software (Zeiss, Jena, Germany). NAFLD activity score was used to assess the severity of changes in the liver. Measurements of the MCPIP-1 immunoexpression in the liver was performed by measuring the optical density in the ImigeJ software and the ZEN software (Zeiss, Jena, Germany). Measurements were made on photos taken under the 40x lens magnification for histological structures such as bile ducts, portal venous and lymphatic vessels, central veins, hepatocytes in zones I and III of hepatic lobules. For each of the indicated structures, 35 measurements were made for each sample.

### Statistical analysis

The results are expressed as the median ± interquartile range (IQR). Nonparametric one-way ANOVA with Tukey’s posttest for multiple comparisons was performed using GraphPad Prism software. For analyses of correlations, the Pearson correlation coefficient was used. The p values are marked with asterisks in the charts (* p<0.05; ** p<0.01; *** p<0.001).

## Results

### Morphological and biochemical characterization of the patients

In the following study, we analyzed a cohort of 36 patients (24 women and 12 men) who underwent operation at the Collegium Medicum Jagiellonian University who were diagnosed according to the European Association for the Study of the Liver. Based on H&E and Oil Red O staining, patients were divided into non-NAFLD (n=5), NAFL (n=12) and NASH (n=19) groups (Fig. 1C, S1, S2). The baseline characteristics of the analyzed groups are summarized in Tables 1 and S2. Blood morphology analysis showed that all patients had elevated levels of white blood cells, monocytes and neutrophils; however, there were no significant differences among the groups (Table S2). Patients in all groups were hyperglycemic, with increased AST and ALT activity (Table 1). In comparison to non-NAFLD group, the NAFL and NASH groups demonstrated increased BMI, urea concentrations and GTTP activity (Table 1).

**Figure 1.**
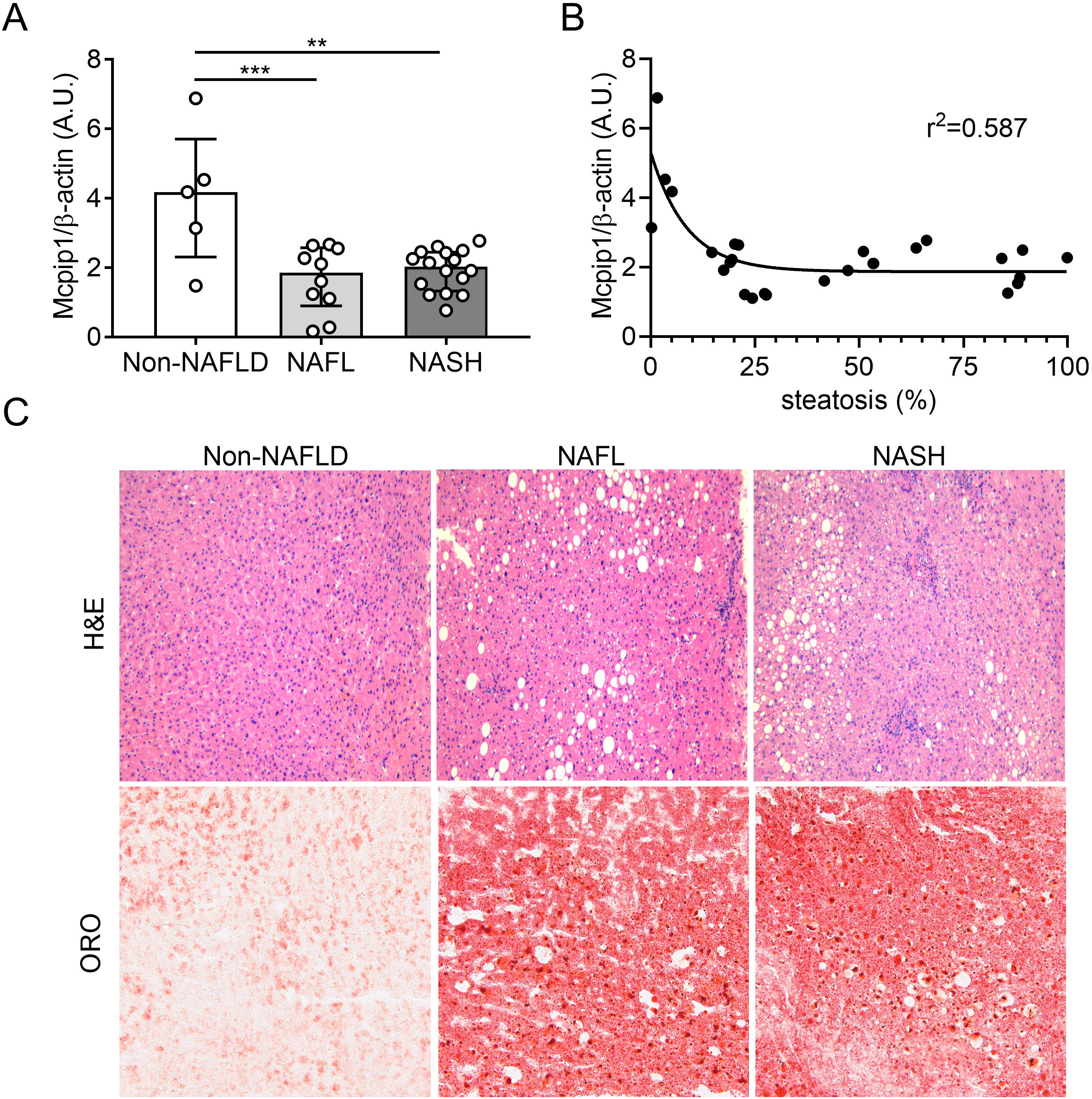
The MCPIP1 level is reduced in the livers of NAFL and NASH patients. A) Densitometric quantification of MCPIP1 protein levels in the livers of control subjects, NAFL patients and NASH patients. B) Correlation between liver MCPIP1 protein levels and % of liver steatosis (% of tissue area stained with lipids). C) Representative H&E and Oil Red O staining of livers from non-NAFLD, NAFL and NASH patients. Statistical significance was determined with Pearson’s correlation. Data were compared using one-way ANOVA with Tukey’s posttest. The graphs show the median ± interquartile range; **p < 0.01; ***p < 0.001.

**Table 1.** Clinical and laboratory characteristic of patients. Data are presented as median (interquartile range in parenthesis) with abnormal values marked in bold and reference values shown in italics. Data were compared using One-way ANOVA with Tukey posttest, * - p < 0.05, ** - p < 0.01 vs. Non-NAFLD group.

|  | Non-NAFLD (n=5) | NAFL (n=12) | NASH (n=19) |
| --- | --- | --- | --- |
| **Age**<br>(years) | 41 (37 – 47) | 41 (35 – 49) | 45 (41 – 51) |
| **Gender**<br>Female/male | 4/1 | 7/5 | 13/6 |
| **BMI**<br>*18.50 – 24.99* | 26.8 (23.8 – 48.2) | 47.4 (40.3 – 54.9) * | 47.0 (39.3 – 51.7) <sup>p=0.09</sup> |
| **Sodium**<br>*136 – 145 mmol/L* | 138 (137 – 140) | 141 (138 – 142) | 141 (139 – 142) |
| **Potassium**<br>*3.5 – 5.1 mmol/L* | 4.17 (3.79 – 4.60) | 4.24 (4.17 – 4.46) | 4.14 (4.03 – 4.47) |
| **Iron**<br>*5.83 – 34.5 μmol/L* | 8.40 (5.43 – 10.30) | 9.27 (7.78 – 10.92) | 8.58 (6.26 – 9.54) |
| **Glucose**<br>*3.3 – 5.6 mmol/L* | **5.65 (5.43 – 8.08)** | **6.10 (5.36 – 6.82)** | **6.41 (5.73 – 8.41)** |
| **HbA1C**<br>*4.3% – 5.9%* | 5.7 (5.3 – 7.3) | 5.7 (5.3 – 6.3) | **6.0 (5.7 – 7.5)** |
| **Bilirubin**<br>*0 - 21 μmol/L* | 7.0 (6.4 – 11.6) | 10.0 (8.4 – 13.5) | 10.4 (8.2 – 14.2) |
| **Urea**<br>*2.76 – 8.07 mmol/L* | 2.8 (2.4 – 3.5) | 5.0 (3.7 – 5.5) * | 3.8 (3.5 – 5.1) * |
| **Creatinine**<br>*44 – 106 μmol/L* | 69 (56 – 86) | 73 (59 – 85) | 74 (65 – 94) |
| **Total protein**<br>*66 – 87 g/L* | 69.2 (60.8 – 74.9) | 70.4 (61.0 – 73.6) | 67.0 (62.3 – 69.6) |
| **Albumins**<br>*35 – 52 g/L* | 41.6 (34.8 – 45.5) | 43.2 (38.1 – 45.2) | 41.9 (37.9 – 43.5) |
| **Cholesterol**<br>*3.2– 5.2 mmol/L* | 4.4 (4.1 – 4.9) | 4.2 (3.9 – 4.9) | 3.8 (3.3 – 4.5) |
| **HDL**<br>*>1.2 mmol/L* | 1.23 (1.09 – 1.73) | **1.19 (0.87 – 1.45)** | **1.16 (1.05 – 1.29)** |
| **LDL**<br>*<3.4 mmol/L* | 2.45 (2.10 – 3.01) | 2.45 (2.26 – 2.74) | 2.37 (1.41 – 2.90) |
| **Triglycerides**<br>*<2.26 mmol/L* | 1.51 (0.85 – 2.18) | 1.31 (1.10 – 2.06) | 1.31 (1.13 – 1.84) |
| **AST**<br>*5 – 40 U/L* | **51 (31 – 104)** | **48 (43 – 94)** | **82 (62 – 112)** |
| **ALT**<br>*5 – 41 U/L* | **62 (33 – 157)** | **65 (47 – 139)** | **91 (78 – 114)** |
| **GTTP**<br>*5 – 61 U/L* | 10 (8 – 18) | 29 (24 – 39) * | 29 (23 – 58) ** |
| **ALP**<br>*35 – 129 U/L* | 61 (55 – 88) | 59 (50 – 65) | 66 (50 – 73) |
| **Cholinesterase**<br>*5320 – 12921 U/L* | 7313 (6463 – 7693) | 7746 (6872 – 8798) | 7604 (6924 – 8746) |
| **Ferritin**<br>*13 – 400 U/L* | 60 (37 – 78) | 232 (88 – 650) * | 125 (87 – 202) |

### Analysis of MCPIP1 liver levels and tissue-specific distribution

The MCPIP1 protein level in the liver was significantly lower in NAFL (MED=1.86, IQR 0.90-2.58) and NASH patients (MED=2.03, IQR 1.33-2.45) than in non-NAFLD control individuals (MED=4.18 IQR 2.31-5.71) (Fig. 1A, S3A). We have previously shown that MCPIP1 is a potent inhibitor of adipogenesis and that its level in adipose tissue inversely correlates with patients’ BMI [11, 12]. Thus, in the next step, we analyzed the amount of MCPIP1 protein in the patients’ livers in relation to hepatic steatosis and BMI. As shown in Figure 1, the MCPIP1 level inversely correlated with hepatic steatosis (r^2^□=□0.587; Fig. 1B) but not with BMI (Fig. S3B). Finally, analysis of cellular MCPIP1 localization revealed that its level was the lowest in the central vein and in the liver parenchyma (both in centrilobular and perilobular hepatocytes) (Fig. 2). Additionally, in all patient groups, the MCPIP1 level was the highest in bile duct epithelial cells (cholangiocytes) and in lymphatic and blood vessels in the portal area (Fig. 2).

**Figure 2.**
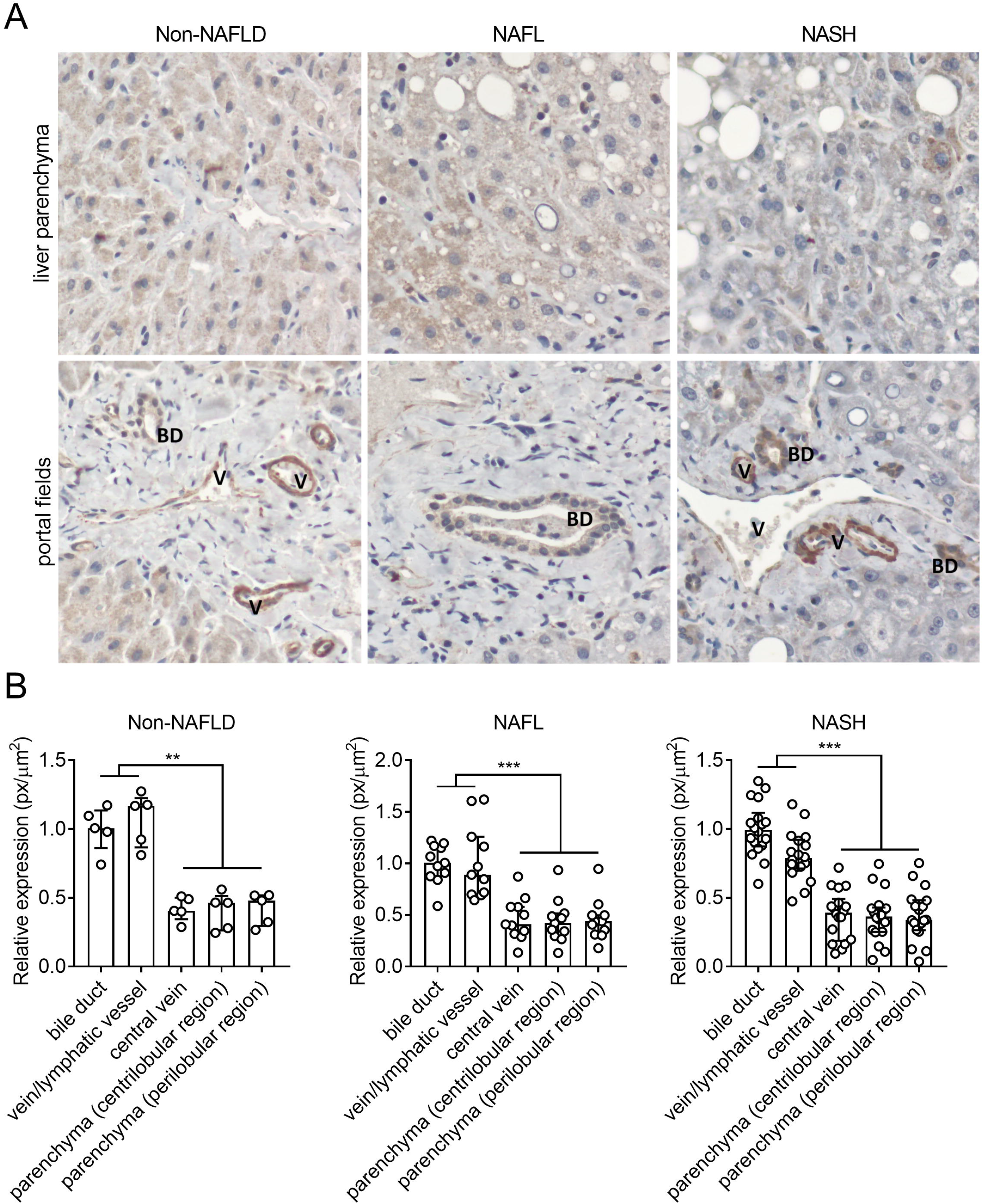
Liver MCPIP1 is predominantly present in bile ducts, veins and lymphatic vessels. A) Representative immunohistochemical staining of MCPIP1 in liver the parenchyma and portal fields. B) Quantitative analysis of MCPIP1 levels in distinct liver areas in non-NAFLD, NAFL and NASH patients. Data were compared using one-way ANOVA with Tukey’s posttest. The graphs show the median ± interquartile range; **p < 0.01; ***p < 0.001.

### Evaluation of insulin signaling and metabolic signaling in the livers of NAFLD patients

The hepatic levels of insulin receptor (IR) were reduced in NAFL patients (MED=0.30, IQR 0.13-0.53) in comparison to the non-NAFLD patients (MED=0.96, IQR 0.36-1.36); however, there was no difference in its phosphorylation (Fig. 3A,B, S4).

**Figure 3.**
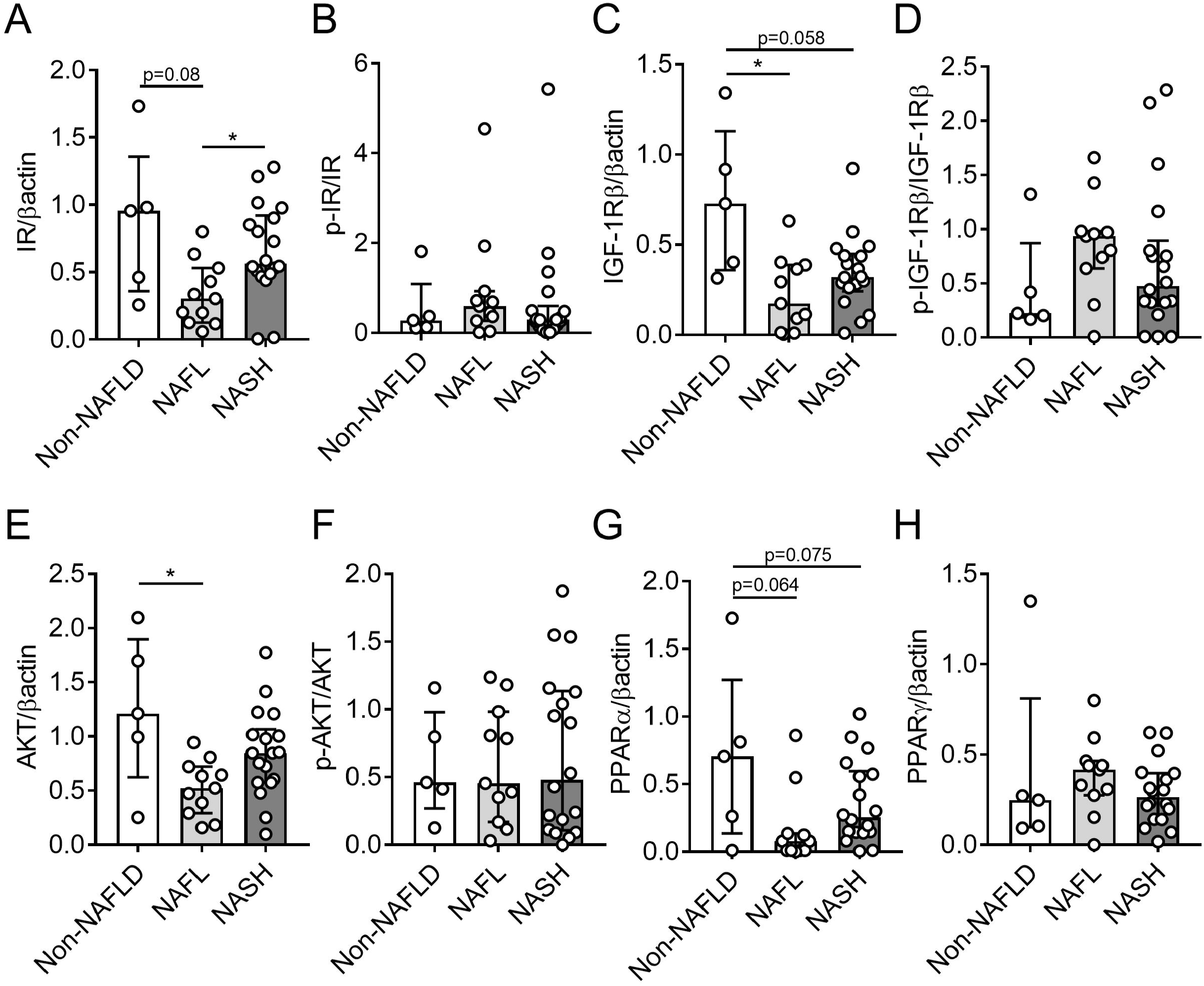
Liver IR, IGF-1Rβ, AKT and PPARα protein levels are decreased in NAFLD patients. Densitometric quantification of A) insulin receptor (IR), B) p-IR, C) insulin-like growth factor 1 receptor beta (IGF-1Rβ), D) p-IGF-1Rβ, E) protein kinase B (AKT), F) p-AKT, G) peroxisome proliferator-activated receptor alpha (PPARα), and H) peroxisome proliferator-activated receptor gamma (PPARγ) protein levels in the livers of control subjects, NAFL patients and NASH patients. Data were compared using one-way ANOVA with Tukey’s posttest. The graphs show the median ± interquartile range; *p < 0.05.

Similarly, the protein level of insulin-like growth factor 1 receptor (IGF-1R) was significantly reduced in the NAFL cohort (MED=0.17, IQR 0.01-0.39) compared with the non-NAFLD cohort (MED=0.73, IQR 0.36-1.13), without differences in its phosphorylation (Fig. 3C,D S4). Additionally, NAFL patients had a diminished amount of protein kinase B (AKT) (MED=0.52, IQR 0.29-0.72) when compared to non-NAFLD patients (MED=1.21, IQR 0.62-1.89), but this was not observed for its phosphorylated form (Fig. 3E, F, S4). Patients from the NASH group had only a reduced amount of IGF-1R levels (MED=0.32, IQR 0.24-0.45) in comparison to non-NAFLD group (MED=0.73, IQR 0.36-1.13) (Fig. 3, S4).

Finally, there was a tendency for higher protein levels of peroxisome proliferator-activated receptor alpha (PPARα) in the livers of non-NAFLD patients (MED=0.71, IQR 0.14-1.27) than in both NAFL (MED=0.08, IQR 0.01-0.14) and NASH patients (MED=0.25, IQR 0.14-0.59) (Fig. 3G, S4). There was no difference in the expression of a second nuclear receptor, namely Peroxisome proliferator-activated receptor gamma (PPARγ) (Fig. 3H, S4).

### Level of expression of MCPIP1 in PBMCs inversely correlates with patient BMI

We have recently demonstrated that the deletion of Mcpip1 in myeloid leukocytes in mice leads to dyslipidemia, low plasma glucose levels and proinflammatory phenotypes that impact NAFLD development [8]. However, in the current study, we showed that the MCPIP1 levels in PBMCs from humans did not differ among non-NAFLD, NAFL and NASH patients (Fig. 4A), but they were negatively correlated with patient BMI (p□=□0.018; r^2^□=□0.198; Fig. 4B,D) and CEBP/β transcript levels (p = 0.025; r^2^ = 0.249; Fig. 4C).

**Figure 4.**
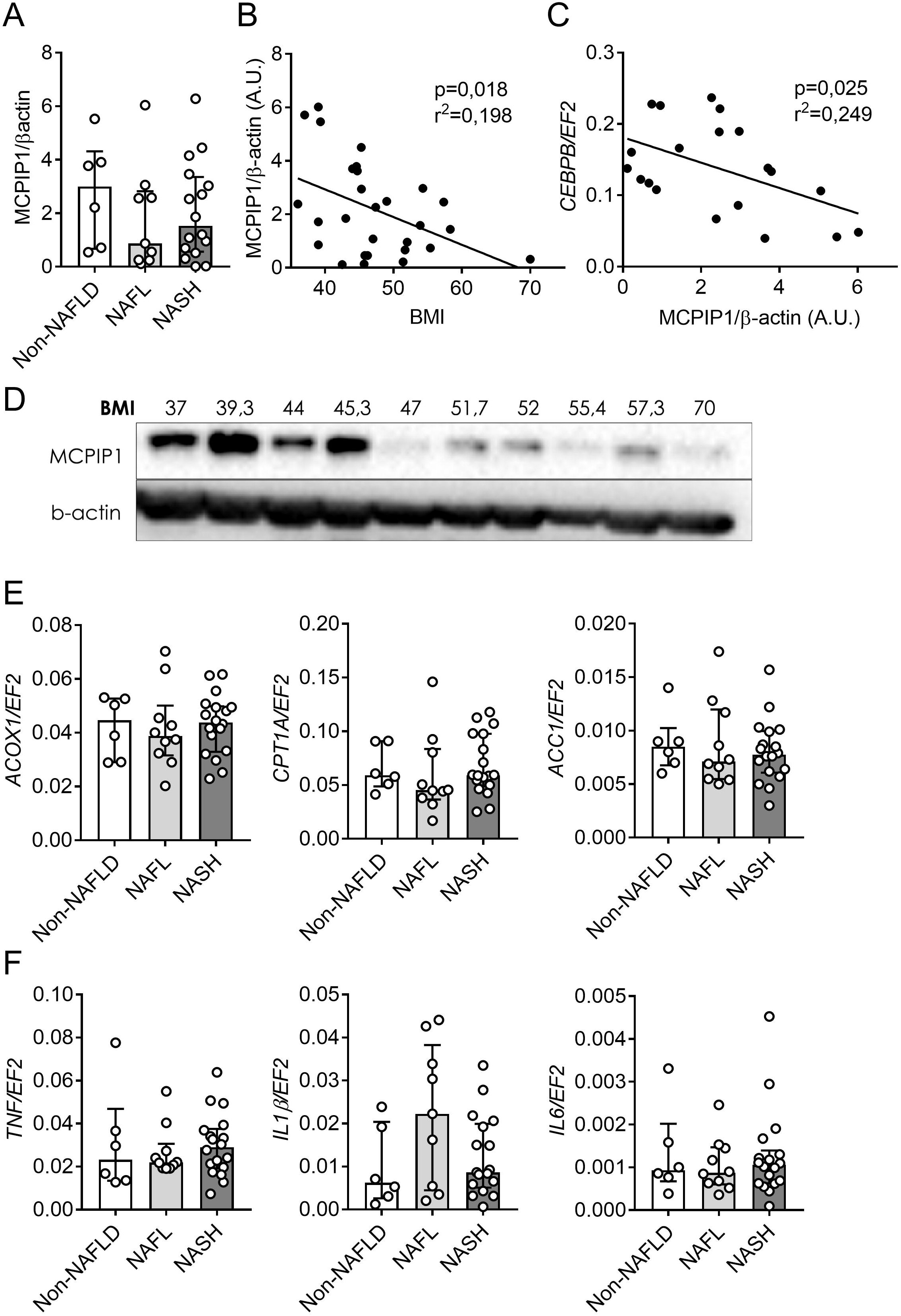
The MCPIP1 level in PBMCs negatively correlates with patient BMI. A) Densitometric quantification of MCPIP1 protein levels in PBMCs of control subjects, NAFL patients and NASH patients. B) Correlation between BMI and MCPIP1 protein levels in PBMC. C) Correlation between MCPIP1 protein levels in PBMCs and C/EBPβ mRNA. D) Representative western blot showing the correlation between MCPIP1 protein levels and patient BMI. Expression of the E) *ACOX1, CPT1A*, and *ACC1* and E) *TNF, IL-1β*, and *IL-6* genes in PBMCs of control subjects, NAFL patients and NASH patients. Data were compared using one-way ANOVA with Tukey’s posttest. Statistical significance was determined with Pearson’s correlation. The graphs show the median ± interquartile range.

Next, we determined the expression levels of key genes that regulate β-oxidation (*ACOX1, CPT1A*, and *ACC1*) and inflammation (*TNF, IL1B, IL6, IL8, IL10*, and *CCL2*) and transcription factors that control metabolism (*FAS, LCN2, CEBPB, SREBP1, PPARA*, and *PPARG*) in PBMCs patients. None of these genes were differentially expressed between non-NAFLD patients and the NAFL or NASH groups (Fig. 4E,F, S5).

## Discussion

In the present study, we have shown that MCPIP1 levels are reduced in liver biopsies collected from NAFLD patients, including both NAFL and NASH patients, in comparison to non-NAFLD control individuals. the possible link between NAFLD and MCPIP1 results from its involvement in lipid metabolism and the regulation of inflammation, as both processes are hallmarks of NAFLD development and progression. MCPIP1 plays an important role in the inhibition of adipogenesis by direct degradation of CCAAT/Enhancer Binding Protein beta (C/EBPβ) transcript [11]. Moreover, ectopic overexpression of MCPIP1 in differentiating mouse preadipocytes impaired adipogenesis not only by the direct cleavage of C/EBPβ mRNA but also by modulating the cellular miRNA pool [13]. Additionally, Mcpip1 levels were lower in primary hepatocytes isolated from high-fat diet-fed mice than in control cells, and thus, it possibly functions to facilitate hepatic lipid accumulation. Additionally, the expression level of Mcpip1 was depleted in visceral fat isolated from obese and glucose-intolerant mice characterized by fatty liver disease in comparison to lean controls [8].

To date, there is only one report describing MCPIP1 levels in patients afflicted with obesity [12]. Losko and coworkers analyzed MCPIP1 expression in biopsies of subcutaneous (SAT) and visceral (VAT) adipose tissue of lean and obese subjects with body mass indices ranging from 27 to 57. In both SAT and VAT, there was a correlation between the MCPIP1 protein level and BMI, as decreased protein levels of MCPIP1 correlated with increased BMI [12]. We found a similar correlation between MCPIP1 levels in PBMCs and BMI, but this correlation was not observed for the expression levels of MCPIP1 in the liver. Additionally, in PBMCs, the level of MCPIP1 protein was inversely correlated with the C/EBPβ transcript, which might be explained by the direct degradation of C/EBPβ mRNA by this RNase [14].

One of the key master regulators of liver metabolism is PPAIRα, which is a ligand-activated transcription factor. PPAIRα regulates the expression of genes involved in fatty acid uptake, beta oxidation, ketogenesis, bile acid synthesis, and triglyceride turnover and thus plays a critical role in the control of metabolism [15]. PPAIRα deficiency leads to excessive lipid accumulation in the liver, resulting in spontaneous steatosis in mice fed a chow diet [16]. Moreover, PPARα deficiency promotes NAFLD and liver inflammation in mice fed a HFD [17]. In humans with NAFLD, hepatic expression of PPARα is decreased, which is in line with our observations [18]. Interestingly, PPAIRα levels were also found to increase in parallel with NAFLD histological improvements secondary to lifestyle intervention or bariatric surgery [15, 18]. Additionally, pharmacological activation of PPAIRα with fenofibrate, gemfibrozil or bezafibrate has beneficial effects in NAFLD patients [19, 20, 21].

MCPIP1 cleaves RNA molecules to regulate a plethora of cellular processes, such as inflammation, cellular differentiation, angiogenesis and adipogenesis [22, 23, 24 11]. By direct degradation of transcripts encoding proinflammatory cytokines (e.g., If-1β, IL-6, and IL-8) [5, 25], MCPIP1 tightly controls the immune system. A constitutive deletion of Mcpip1 in mice leads to a lethal phenotype resulting from severe systemic and multiorgan inflammation [7]. Mcpip1 deletion in mice was also shown to trigger autoimmune diseases, such as gastritis (whole-body Mcpip1 knockout) [26], lupus (deletion in myeloid leukocytes) [27], and primary biliary cholangitis (deletion in liver epithelial cells) [9].

Our results show the cellular distribution of the MCPIP1 protein in human livers and highlight its important role in the biology of cholangiocytes and endothelial cells, where the MCPIP1 level was the highest in all experimental groups. In line with these immunohistochemistry data, both the Human Protein Atlas and Liver Single Cell Atlas demonstrate cholangiocytes as a population with the highest *ZC3H12A* expression in comparison to other liver cells [(https://www.proteinatlas.org/,28]. The Liver Single Cell Atlas was generated by Brancale and Vilarinho, who integrated and analyzed available human liver single-cell RNA-sequencing (scRNA-seq) datasets. The authors used results from gene expression data across a variety of annotated parenchymal and nonparenchymal cells derived from 28 healthy human livers analyzed by 5 independent studies. In addition to cholangiocytes, a high level of *ZC3H12A* was also detected in lymphoid cells and in hepatocytes [28]. Finally, Mcpip1 deficiency in murine liver epithelial cells leads to the development of PBC symptoms. Thus, it would be interesting to analyze *ZC3H12A* expression in livers from PBC patients and tumors collected from cholangiocarcinoma and hepatocellular carcinoma biopsies.

Although liver biopsy is still the gold standard for NAFLD assessment, a less invasive collection of liquid biopsy involving blood sampling is highly appreciated. Thus, a liquid biopsy containing peripheral blood mononuclear cells, extracellular vehicles or circulating DNA might be used as a diagnostic and monitoring tool. PBMCs are widely used as screening materials for the identification of new disease-associated biomarkers. They can reflect the gene expression profile involved in a number of pathological conditions, including obesity, inflammation or oxidative stress [29, 30]. Importantly, they express receptors for insulin, glucagon and leptin on their surface, and thus they can respond to hormonal changes that reflect the metabolic response of various organs [31, 32]. Moreover, the human MCPIP1 transcript was originally shown to be highly expressed in leukocytes [33]. The PBMC expression profile might also be used for discriminating NASH from NAFL patients because the expression levels of cytokine- and chemokine-encoding genes (IFNγ, IL-2, IL-15, CCL2, CXCL11, and IL-10) in PBMCs were significantly upregulated in PBMCs from NASH patients in comparison to those from NAFL patients [34]. Although the expression of genes regulating β-oxidation and inflammation and controlling metabolism selected by us did not differ in these cohorts of NAFL and NASH patients, Kado and coworkers were able to successfully divide most of 54 NAFLD patients into NAFL and NASH groups based on INFγ, CCL2 and IL-10 expression levels [34]. It is also possible to use flow cytometry analysis of blood leukocytes to discriminate NAFL and NASH patients. Both peripheral lymphocyte and myeloid cell subsets differ in terms of their abundance, activation, polarization or cell membrane markers [35, 36].

Taken together, our data demonstrate that the MCPIP1 protein level is reduced in the livers of NAFL and NASH patients and is predominantly expressed in cholangiocytes, veins and lymphatic vessels. Our data also revealed that the amount of PPARα was reduced in NAFLD patients, which is in agreement with recent studies. Although the MCPIP1 level in PBMCs was not affected by NAFLD, its level correlated with patients’ BMI and C/EBPβ transcript level. Further research is required to investigate the specific role of MCPIP1 in both NAFL initiation and the transition to NASH. Such experiments might be performed, for example, on tissue-specific Mcpip1 knockout mouse models subjected to diet-induced obesity or sterile inflammation.

## Abbreviations

ACC1: Acetyl-CoA carboxylase 1
ACOX1: Acyl-CoA Oxidase 1
AKT: Protein kinase B
ALT: alanine aminotransferase
AST: aspartate aminotransferase
BMI: body mass index
C/EBPβ: CCAAT/enhancer-binding protein
CCL2: C-C Motif Chemokine Ligand 2
CPT1a: Carnitine Palmitoyltransferase 1A
CXCL11: C-X-C motif chemokine 11
EDTA: ethylenediaminetetraacetic acid
FAS: Fatty Acid Synthase
GTTP: Gamma-glutamyl Transpeptidase
HBV: hepatitis virus B
HCV: hepatitis virus C
HFD: high fat diet
HIV: human immunodeficiency virus
IFNγ: interferon gamma
IGF-1R: Insulin-like Growth Factor 1 Receptor
IL: interleukin
IQR: interquartile range
IR: Insulin Receptor
LCN2: Lipocalin-2
MCPIP1: Monocyte Chemotactic Protein-1-Induced Protein-1
MED: median
NAFLD: Non-alcoholic fatty liver disease
NAFL: Non-alcoholic fatty liver
NASH: Non-alcoholic steatohepatits
PBMCs: Peripheral blood mononuclear cells
PPAR: Peroxisome proliferator-activated receptor
SAT: subcutaneous adipose tissue
SREBP1: Sterol Regulatory Element-binding protein 1
TNF: Tumor Necrosis Factor
VAT: visceral adipose tissue

## Acknowledgment

We thank Agnieszka Jasztal, Edyta Kus and Stefan Chlopicki from Jagiellonian Centre for Experimental Therapeutics at Jagiellonian University for excellent help with liver histology staining.

