## Supplementary figures and images for "Hepatic MCPIP1 protein levels are reduced in NAFLD patients and are predominantly expressed in cholangiocytes and liver endothelium"

### Supplemental Figure 1

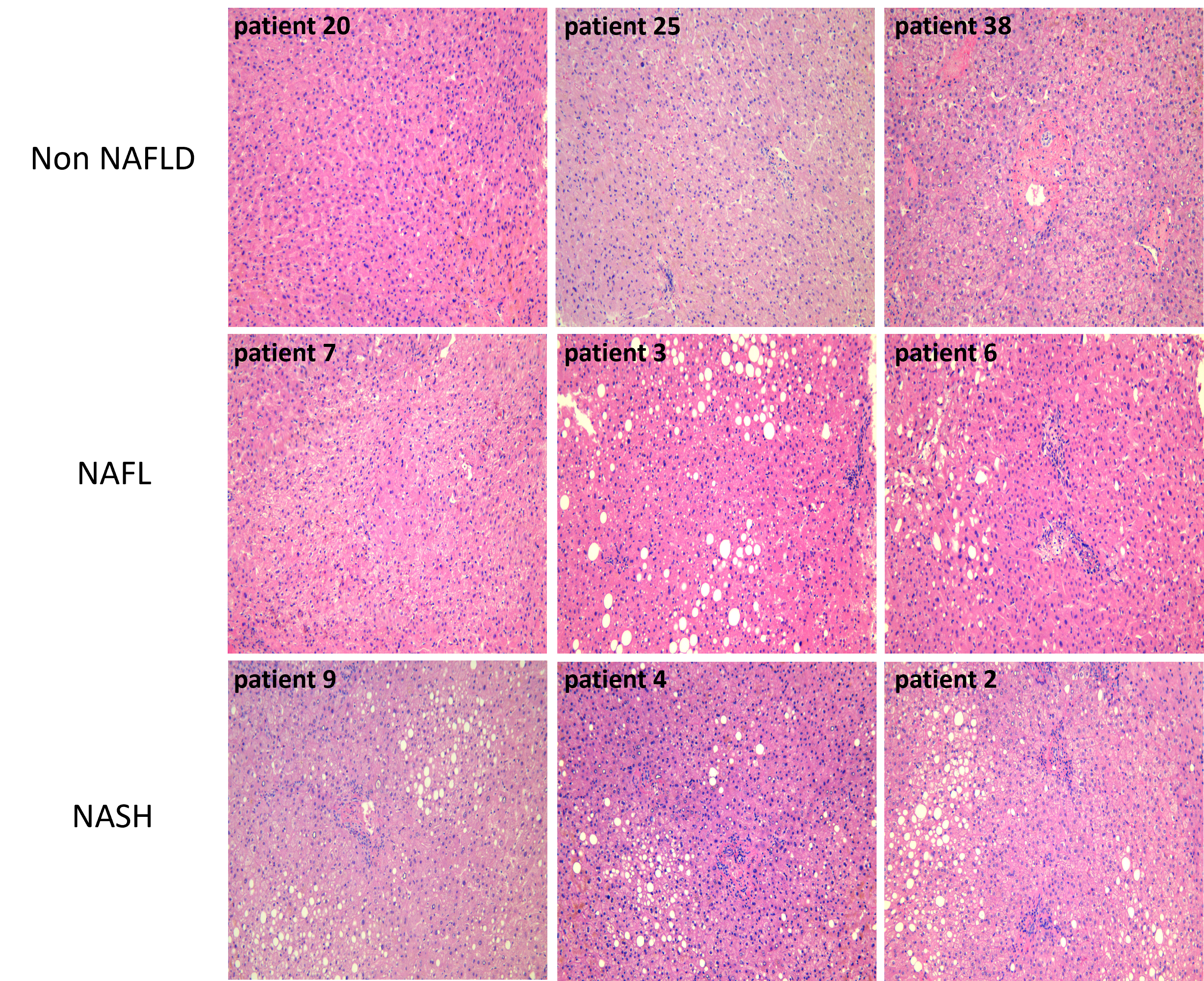

### Supplemental Figure 2

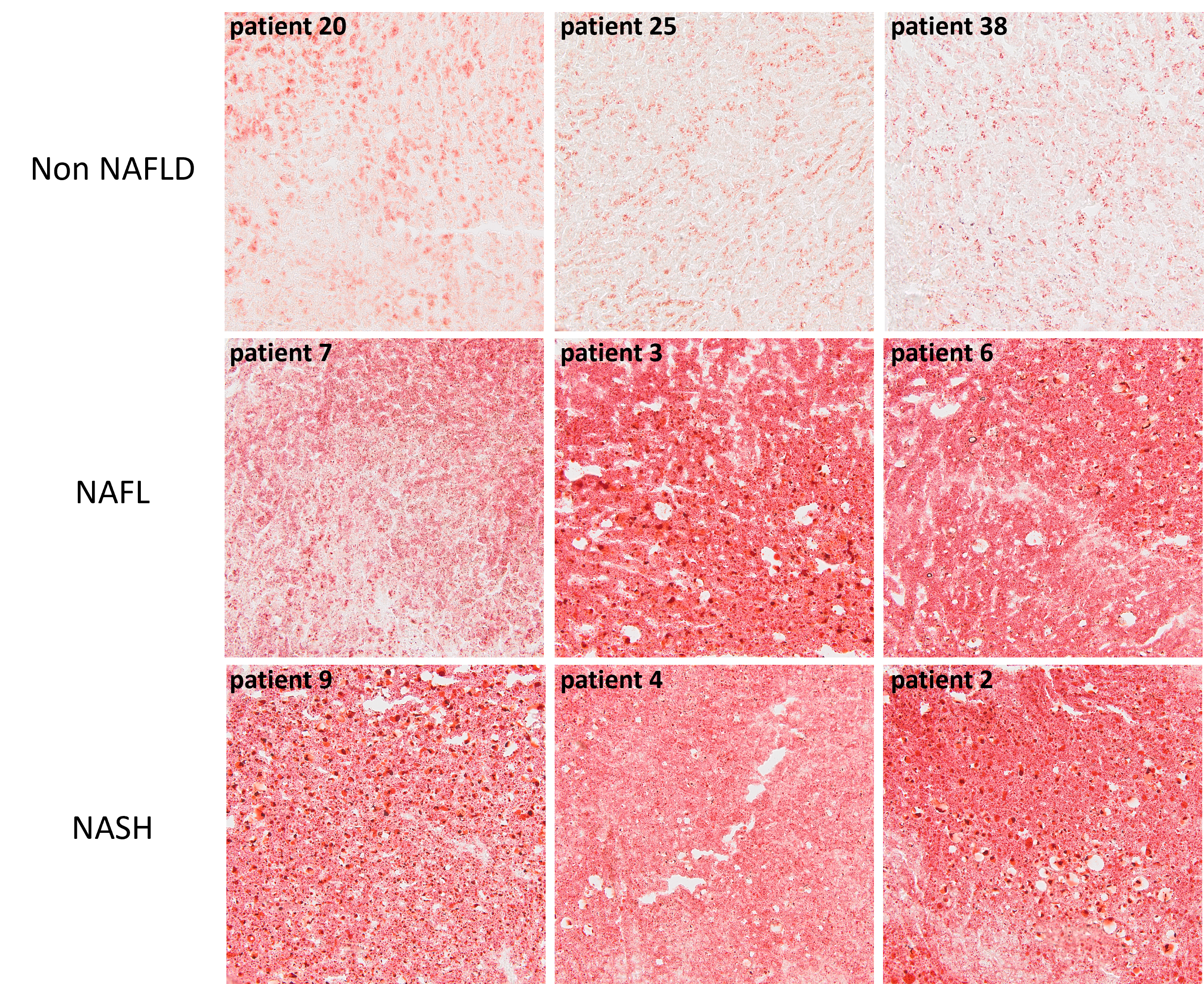

### Supplemental Figure 3

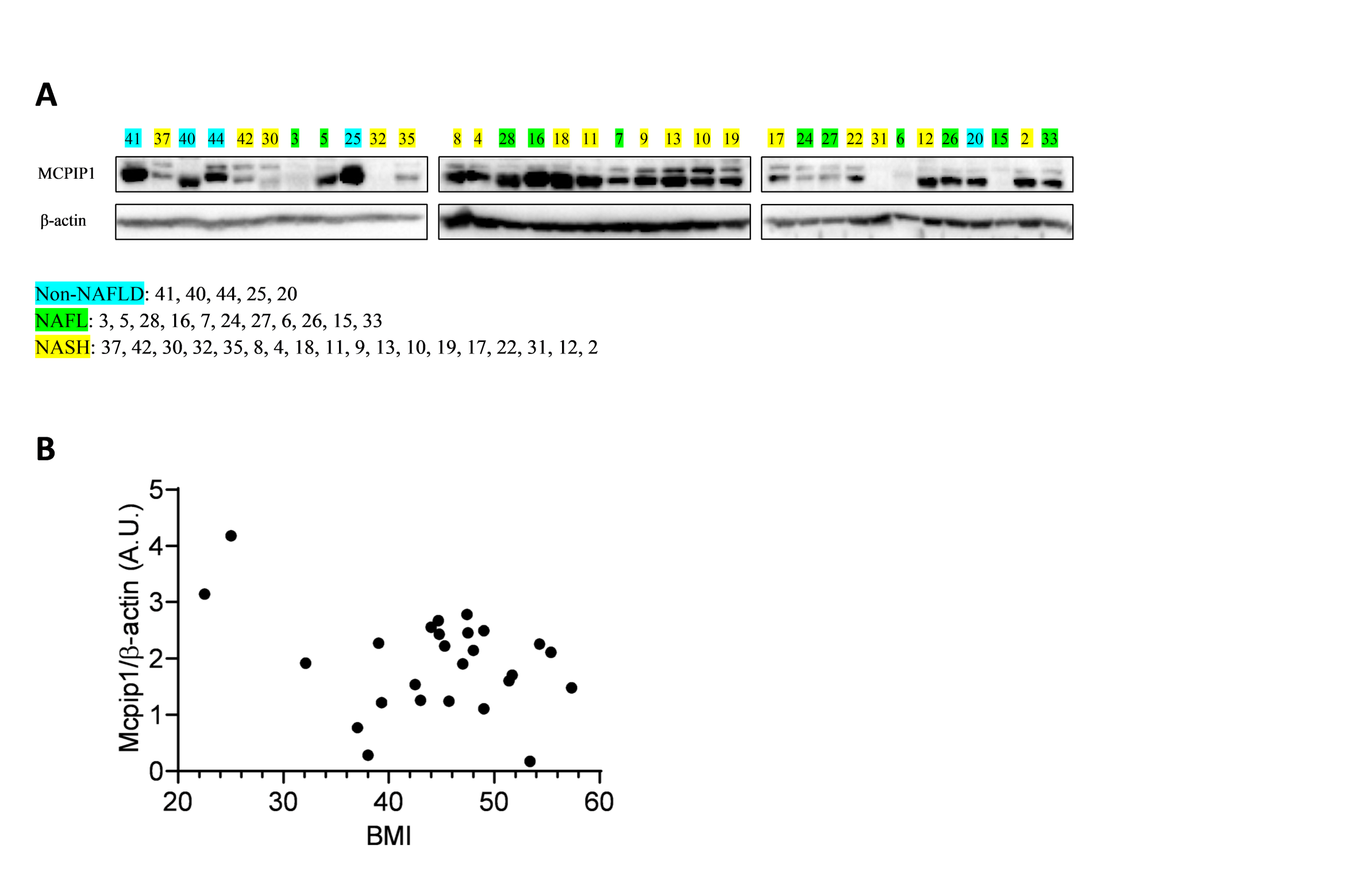

### Supplemental Figure 4

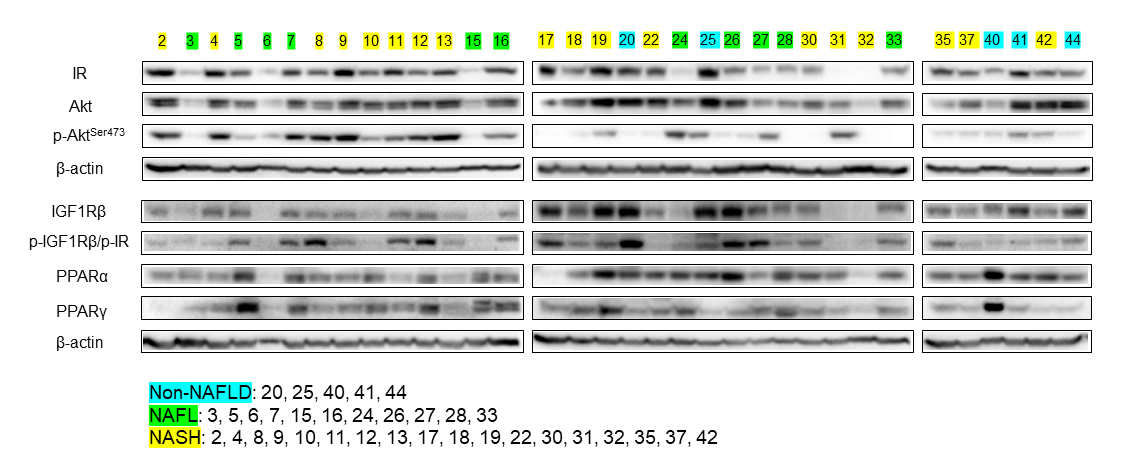

### Supplemental Figure 5

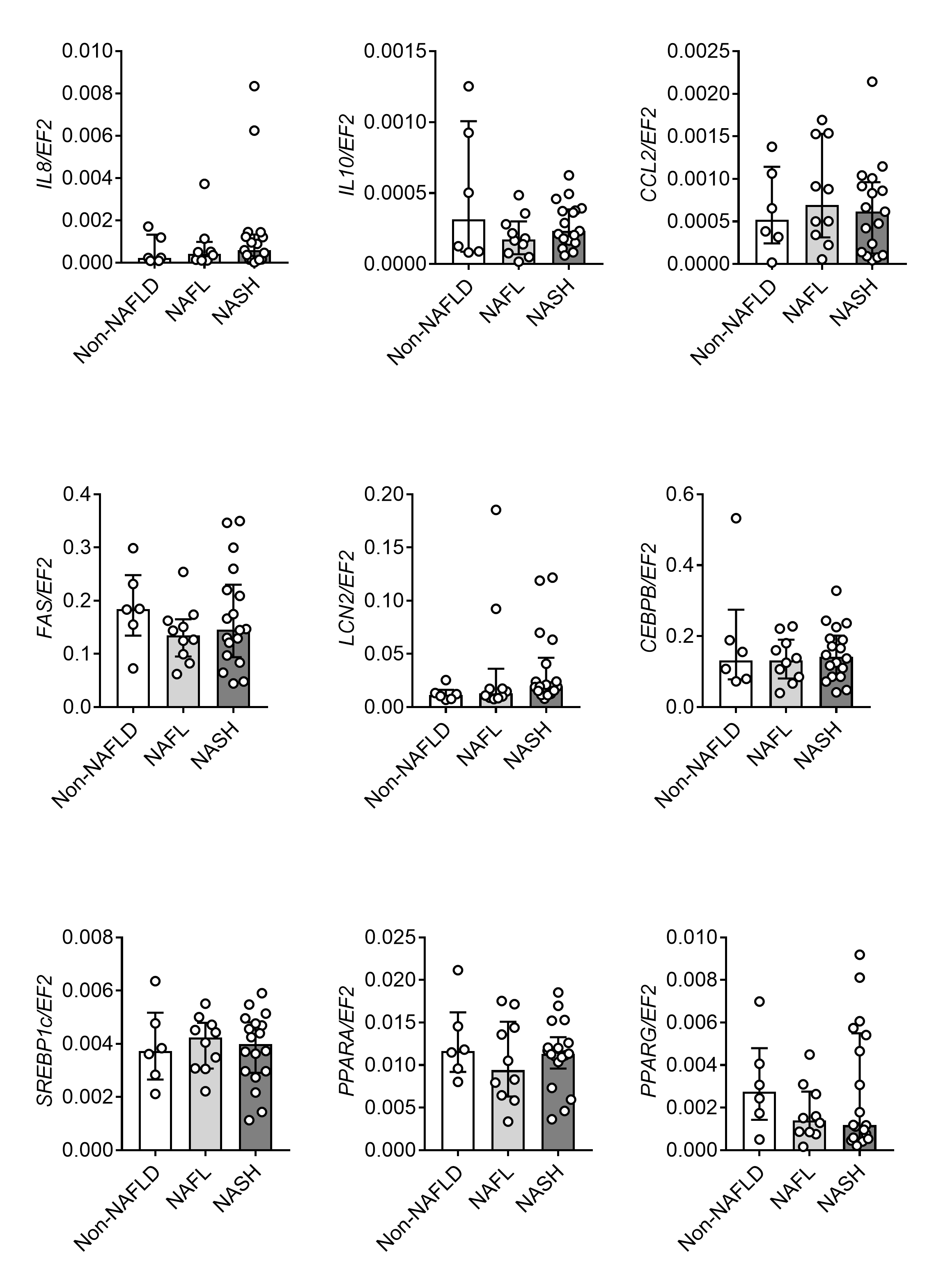
