## Supplemental Figure Legends for "Hepatic MCPIP1 protein levels are reduced in NAFLD patients and are predominantly expressed in cholangiocytes and liver endothelium"

**Figure S1.** H&E staining of livers from non-NAFLD, NAFL and NASH patients.

**Figure S2.** ORO staining of livers from non-NAFLD, NAFL and NASH patients.

**Figure S3.** A) MCPIP1 protein level in livers of all analyzed patients. B) Dot plot showing patients’ BMI vs. liver MCPIP1 protein level.

**Figure S4.** Western blot analysis of proteins in the livers of all analyzed patients. Insulin receptor (IR); protein kinase B (AKT); insulin-like growth factor 1 receptor beta (IGF-1Rβ); peroxisome proliferator-activated receptor alpha (PPARα); peroxisome proliferator-activated receptor gamma (PPARγ).

**Figure S5.**

Expression of genes coding for proteins regulating inflammation (*IL8, IL10, CCL2,* and *LCN2*), lipid metabolism (*FAS*), and transcription factors controlling metabolism (*CEBPB, SREBP1, PPARA,* and *PPARG*) in PBMCs of control subjects, NAFL patients and NASH patients. Data were compared using one-way ANOVA with Tukey’s posttest. The graphs show the median ± interquartile range.
