## Supplemental Tables for "Hepatic MCPIP1 protein levels are reduced in NAFLD patients and are predominantly expressed in cholangiocytes and liver endothelium"

**Table S1. Sequences of primers used in the study.**

| Gene | Sequence |
| --- | --- |
| *EF2* | F: GACATCACCAAGGGTGTGCAG  R: TTCAGCACACTGGCATAGAGGC |
| *ACOX1* | F: CTTTGCCAGGAATTACCGTTGG  R: GGTCCCGTAAGTCAGCTTGT |
| *CPT1a* | F: TCCAGTTGGCTTATCGTGGTG  R: CTAACGAGGGGTCGATCTTGG |
| *ACC1* | F: CTCTCACGCTCAAGTCACCA  R: GCAATAAGAACCTGGCGTGC |
| *TNF* | F: CTTCTCGAACCCCGAGTGAC  R: ATGAGGTACAGGCCCTCTGA |
| *IL-1β* | F: TGGGTAATTTTTGGGATCTACACTCT  R: AATCTGTACCTGTCCTGCGTGTT |
| *IL-6* | F: GTGAAAGCAGCAAAGAGGCA  R: TCACCAGGCAAGTCTCCTCA |
| *IL-8* | F: ATGACTTCCAAGCTGGCCGTGGCT  R: TCTCAGCCCTCTTCAAAAACTTCTC |
| *IL-10* | F: CAAGGCGCATGTGAACTCC  R: GATGCCTTTCTCTTGGAGCTTATT |
| *CCL2* | F: CTTCTGTGCCTGCTGCTCATAGC  R: CCAGGTGGTCCATGGAATCCTG |
| *FAS* | F: GGACAGAGCAACTACGGCTT  R: GTGCTCATCGTCTCCACCAA |
| *LCN* | F: TGTCACCTCCGTCCTGTTTAG  R: TCTCCCGTAGAGGGTGATCTT |
| *CEBPB* | F: AGCGACGAGTACAAGATCCG  R: GCTGCTCCACCTTCTTCTGC |
| *SREBF1* | F: CCATGGATTGCACTTTCGAA  R: GGCCAGGGAAGTCACTGTCTT |
| *PPARA* | F: ATGAGGCCATATTCGCCATGC  R: GTTGCTCTGCAGGTGGAGTCT |
| *PPARG* | F: AGTGGAGACCGCCCAGGTTTGCT  R: CCTGCAGTAGCTGCACGTGTTCCGT |

**Table S2. Blood count.** Data are presented as median (interquartile range in parenthesis) with abnormal values marked in bold and reference values shown in italics. Data were compared using One-way ANOVA with Tukey posttest, * - p < 0.05, ** - p < 0.01 vs. Non-NAFLD group.

|  | **Non-NAFLD (n=5)** | **NAFL (n=12)** | **NASH (n=19)** |
| --- | --- | --- | --- |
| **White blood cells**  *4 – 10 * 10^3^/µL* | **13.20 (12.72 – 15.05)** | **12.42 (10.88 – 15.56)** | **10.75 (8.80 – 12.06)** |
| **Lymphocytes**  *0.8 – 4 * 10^3^/µL* | 2.0 (1.3 – 2.6) | 1.6 (1.2 – 2.2) | 1.5 (1.3 – 1.6) |
| **Lymphocytes**  *20 – 40 %* | **16.1 (9.4 – 19.3)** | **13.4 (9.9 – 18.3)** | **14.9 (11.7 – 18.4)** |
| **Monocytes**  *0.16 – 0.8 * 10^3^/µL* | **0.9 (0.7 – 1.1)** | **1.2 (0.9 – 1.4)** | 0.7 (0.7 – 0.9) |
| **Monocytes**  *4 – 8 %* | 7.0 (4.8 – 8.1) | **9.3 (8.2 – 10.6)** | 8.0 (6.7 – 8.6) |
| **Neutrophils**  *2.4 – 7 * 10^3^/µL* | **10.5 (9.2 – 12.5)** | **9.7 (8.8 – 12.2)** | **7.5 (6.4 – 9.3)** |
| **Neutrophils**  *58 – 66 %* | **75.8 (71.9 – 84.5)** | **78.9 (70.8 – 81.2)** | **75.9 (71.8 – 80.8)** |
| **Red blood cells**  *3.5 – 6.5 * 10^6^/µL* | 4.36 (4.20 – 5.02) | 4.57 (4.29 – 4.95) | 4.50 (4.15 – 4.62) |
| **Hemoglobin**  *11 – 17 g/dL* | 12.5 (11.8 – 14.0) | 13.5 (12.3 – 14.6) | 13.1 (12.0 – 13.4) |
| **Hematocrit**  *37 – 54 %* | 38.2 (35.0 – 41.7) | 40.1 (36.9 – 43.1) | 38.8 (36.3 – 39.9) |
| **MCV**  *82 – 92 fl* | 84.1 (81.1 – 86.9) | 86.0 (83.8 – 87.3) | 85.8 (83.7 – 91.6) |
| **MCH**  *27 – 31 pg* | 28.0 (27.0 – 29.1) | 29.0 (28.1 – 30.1) | 28.8 (27.6 – 30.0) |
| **MCHC**  *32 – 36 g/dL* | 33.3 (32.8 – 33.9) | 33.6 (32.7 – 34.1) | 33.5 (32.3 – 34.0) |
| **RDW-SD**  *37.3 – 46.7 fl* | 40.8 (39.2 – 42.3) | 40.8 (40.1 – 43.4) | 42.1 (40.8 – 46.5) |
| **RDW-CV**  *12.1 – 14.1 %* | 12.9 (12.8 – 14.0) | 13.3 (12.7 – 13.7) | 13.3 (12.8 – 14.9) |
| **PLT**  *125 – 340 * 10^3^/µL* | 226 (206 – 303) | 235 (222 – 267) | 234 (186 – 277) |
| **MPV**  *9.4 – 12.6 fl* | 11.9 (10.5 – 12.7) | 10.6 (10.0 – 11.8) | 10.9 (10.4 – 11.2) |
| **PCT**  *0.16 – 0.34 %* | 0.24 (0.23 – 0.38) | 0.26 (0.24 – 0.28) | 0.24 (0.21 – 0.28) |
| **PDW**  *9.2 – 16.8 fl* | 14.8 (12.3 – 17.4) | 12.5 (11.3 – 15.1) | 13.3 (12.8 – 14.1) |
| **P-LCR**  *16 – 46.4 %* | 41.2 (29.2 – 45.9) | 29.3 (25.4 – 39.6) | 32.2 (29.2 – 35.2) |
| **Prothrombin time**  *sec* | 11.2 (10.5 – 11.4) | 11.6 (11.1 – 12.2) | 11.6 (11.3 – 11.9) |
| **INR**  *0.9 – 1.2 sec* | 0.97 (0.92 – 1.01) | 1.01 (0.95 – 1.05) | 1.00 (0.96 – 1.03) |
| **APTT**  *26 – 36 sec* | 25.3 (23.9 – 29.9) | 25.9 (24.9 – 27.5) | 28.2 (27.0 – 29.3) |
